# UMI-Reducer: Collapsing duplicate sequencing reads via Unique Molecular Identifiers

**DOI:** 10.1101/103267

**Authors:** Serghei Mangul, Sarah Van Driesche, Lana S. Martin, Kelsey C. Martin, Eleazar Eskin

## Abstract

**Summary:** Every sequencing library contains duplicate reads. While many duplicates arise during polymerase chain reaction (PCR), some duplicates derive from multiple identical fragments of mRNA present in the original lysate (termed “biological duplicates”). Unique Molecular Identifiers (UMIs) are random oligonucleotide sequences that allow differentiation between technical and biological duplicates. Here we report the development of UMI-Reducer, a new computational tool for processing and differentiating PCR duplicates from biological duplicates. UMI-Reducer uses UMIs and the mapping position of the read to identify and collapse reads that are technical duplicates. Remaining true biological reads are further used for bias-free estimate of mRNA abundance in the original lysate. This strategy is of particular use for libraries made from low amounts of starting material, which typically require additional cycles of PCR and therefore are most prone to PCR duplicate bias.

**Availability and Implementation:** The UMI-Reducer is an open source Python software and is freely available for non-commercial use (GPL-3.0) at **https://sergheimangul.wordpress.com/umi-reducer/**. Documentation and tutorials are available at **https://github.com/smangul1/UMI-Reducer/wiki/**.

**Supplementary information:** Flowchart of Library Construction

## 1. Introduction

High throughput RNA sequencing technologies have provided invaluable research opportunities across multiple scientific domains by producing short quantitative readouts (reads) of various types of RNAs present in cells or tissues. Traditionally, the number of mRNA copies (abundance) in the sample is estimated based on number of reads (read counts). Sequencing libraries made using PCR amplification invariably contain duplicate reads. Many duplicates are of technical origin and arise during PCR; however, some duplicates derive from multiple identical fragments of mRNA present in the original lysate (termed "biological duplicates"). The PCR step, which takes place prior to sequencing, can confound accurate transcriptome quantification because PCR amplification is biased according to fragment abundance, fragment length, and GC content. Removing all duplicates based on their mapped location can eliminate the effects of PCR bias. However, this approach to removing technical duplicates will also remove biological duplicates and is not ideal for libraries with low amounts of starting material. Low-input libraries are subject to greater bias, because they require more rounds of PCR. Thus, the best estimate of mRNA abundance in the original lysate is obtained by collapsing PCR duplicates while keeping biological duplicates. Keeping biological duplicates is also advantageous because low-input libraries tend to have lower overall sequencing depth, and thus total unique reads are maximized when biological duplicates are kept.

We developed **UMI-Reducer**, a new computational tool that differentiates PCR duplicates from biological duplicates. **UMI-Reducer** uses 7mer degenerate sequences called Unique Molecular Identifiers (UMIs), which are attached to sequencing adapters prior to PCR amplification^1^. Reads with identical UMIs and sequencing positions are considered technical duplicates and are collapsed to a single unique read. Reads from the same sequencing positions but non-identical UMIs (biological duplicates) are considered as separate unique reads. The final set of all unique reads is used for an accurate estimation of mRNA abundances. The proposed method then annotates the final set of reads to genomic regions (e.g., coding sequence, 5’UTR, 3’UTR, intron, intergenic) and counts the number of reads per gene in each genomic region. For example, in a ribosome profiling library, a user can obtain the total number of coding sequence reads for each gene.

## 2. Methods

### 2.1 Anatomy of a read

We used the Illumina MiSeq protocol to generate 150nt reads. **UMI-Reducer** assumes the structure of a read is composed of the following parts: barcode - a specified 4nt sequence used to uniquely tag the individual sample prior to sequencing; **Unique Molecular Identifiers** (UMI) – random oligonucleotide sequences allowing differentiation between technical and biological duplicates; 3nt degenerate sequence; mRNA fragment to be sequenced (hereafter referred to as “read”); L32 RNA linker of specified sequence^2^; and Illumina adaptor sequence (Figure 1a). The primer associated with each read contains a 7mer degenerate sequence called the Unique Molecular Identifier (UMI), such that each primer is in fact 16,384 uniquely identifiable primers. The degenerate sequence (DDD) represents the amino acids A, G, or T (but not C) and is designed to reduce ligation bias during library preparation. The 20nt 3’ L32 RNA linker sequence is defined, and we use it to determine the 3’ boundary of the sequenced RNA, which is usually of variable length. Reads that do not contain a L32 RNA linker are considered background reads not derived from a linker ligated-product and are discarded. Complete details of library preparation, including all oligos used, are provided in the Supplementary Text.

**Figure 1.**
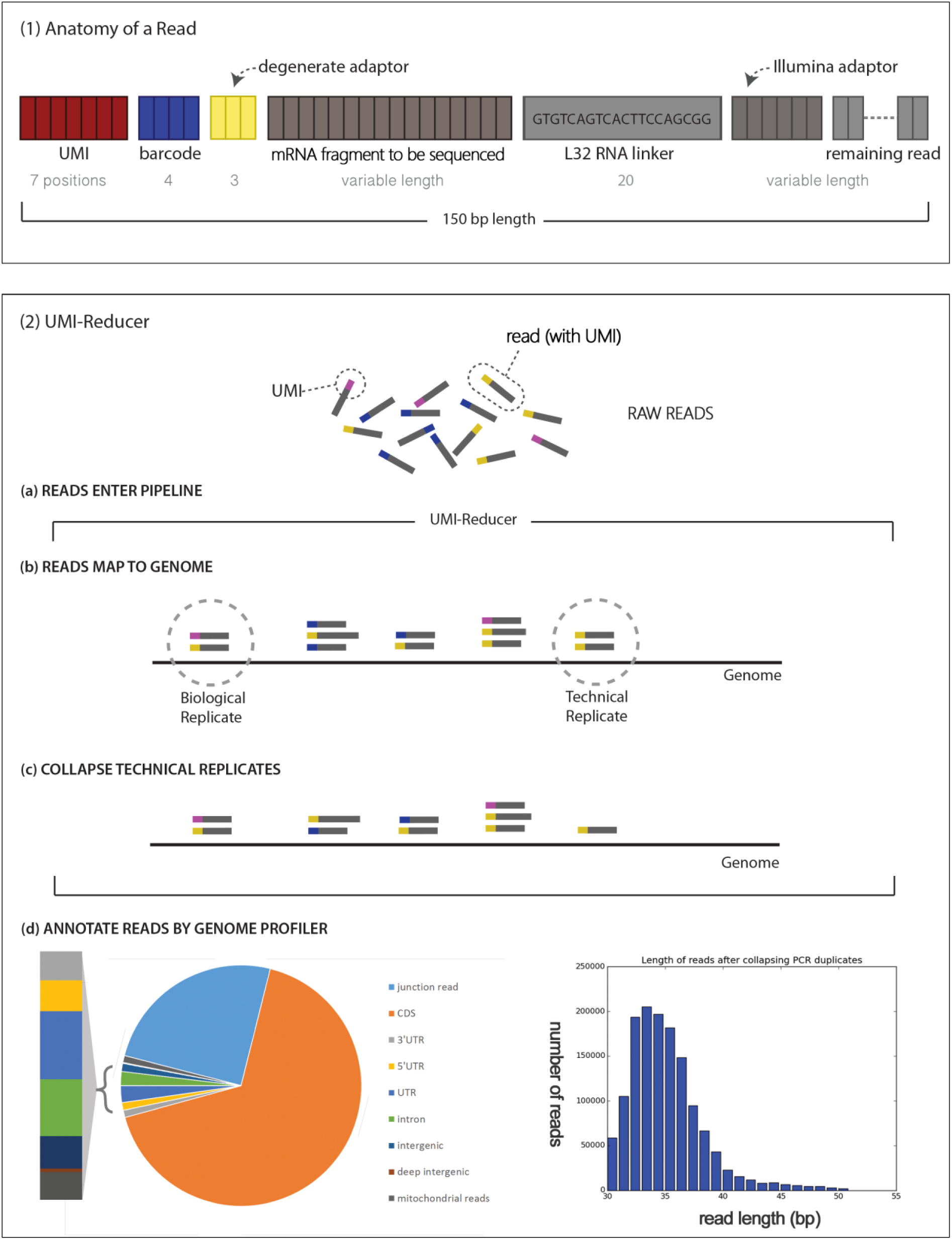
Overview of the UMI-Reducer. **(1)** A schematic of 150 bp sequencing read, obtained by Illumina MiSeq Illumina V3 150 cycle kit. Read is comprised of the following parts: barcode (blue color, 4bp), Unique Molecular Identifiers (UMI) (red color, 7 bp), degenerate sequence (yellow color, 3bp), mRNA fragment to be sequenced (hereafter referred to as “read”) (green color, variable length), L32 RNA linker (grey color, GTGTCAGTCACTTCCAGCGG, 20bp), and Illumina adaptor (grey color, CCGCTGGAAGTGACTGACAC). The degenerate sequence represents the amino acids A, G, or T (but not C) in a degenerate region of three positions. This sequence is designed to reduce ligation bias during library preparation, as the CircLigase enzyme does not prefer the base C. **(2)** UMI-Reducer steps. Raw sequencing data prepared as described in (1) enters the pipeline (2a). Reads containing the full sequence of L32 RNA linker are identified, and the RNA sequence is extracted and saved into a FASTQ file. Corresponding UMI sequences are added to the read ID and used in downstream analyses. Corresponding barcode is used to partition reads into individual samples (not shown). Reads are then mapped to the genome using tophat2 (2b). Reads with matching UMIs that are located in identical positions along the genome (i.e., identical start and end position of the read) are considered technical duplicates; reads with different UMIs that are located in the same position are considered biological duplicates. UMI-Reducer collapses technical duplicates by combining all copies into one read (2c). UMI-Reducer reports, as a pie chart, the genomic profiles of mapped reads after filtering out multi-mapped reads and collapsing PCR duplicates. The genome profiler annotates reads into the following categories (2d): junction reads, CDS, 5’UTR, 3’UTR, intron, intergenic, deep-intergenic (reads mapped 1Kb away from the gene boundaries), and mitochondrial reads (MT). UMI-Reducer reports, as a histogram, length distribution of uniquely mapped reads after collapsing technical duplicates.

### 2.2 Extract Reads

Given raw reads as described in Section 2.2, the *extractTag.py* tool uses barcode sequences to partition reads into individual samples. This tool removes the L32 linker sequence on the 3’ end of the read. Reads that do not contain the full L32 sequence are discarded. Remaining reads are stored in FASTQ format (Figure1b). Corresponding UMI sequences are added to the read ID and used in downstream analysis.

### 2.3 Collapse technical duplicates

The *collapsePCRduplicates.py* is a tool used to collapse PCR duplicates while retaining biological duplicates for more accurate estimation of RNA abundance in the original lysate. We first map the reads on the reference genome using tophat2^3^. Tophat2 produces a bam-formatted alignment file. By default, *collapsePCRduplicates.py* discards the reads that map to more than one locus (multi-mapped reads). Using –m option of *collapsePCRduplicates.py* and *gprofilePlus.py* (ROP v1.0.6)^4^, one can randomly assign multi-mapped reads based on their transcript abundances. The BAM file is parsed by *collapsePCRduplicates.py* to identify reads mapped to the same position on the reference genome (i.e., reads with the same starting and ending position). Reads that are mapped to the same position in the genome and have identical UMIs are categorized as PCR duplicates. These duplicate reads are then collapsed into a single read (Figure1b). The **UMI-Reducer** protocol produces for downstream analysis a BAM file that is comprised of biologically unique reads and their corresponding mapping positions. Additionally, *collapsePCRduplicates.py* produces a series of histograms that show the length distribution of mapped reads before and after collapsing PCR duplicates. These histograms, available as PNG files, can be used to check that the reads are of anticipated length (e.g., for ribosome profiling in mammalian cells we expect the read length to be approximately 30-40nt). The number of reads at the *collapsePCRduplicates.py* step is reported in a STAT file.

### 2.4 Annotate Reads using Genome Profiler

The ROP protocol^4^’s *gprofile.py* annotates each read to genomic region (e.g., coding sequence, 5’UTR, 3’UTR, intron, intergenic) and outputs annotation descriptions on three levels: the number of reads annotated to each genomic feature across the entire sample, the number of reads annotated to each genomic feature across each gene, and annotation for each individual read in the sample. We provide the option to assign multi-mapped reads into genomic features based on the genes’ expression levels, which are determined from uniquely mapped reads. To take advantage of this option, one needs to select the –multi option of *gprofile.py*, which will assign a genomic feature for each mapping of the multi-mapped read. *gprofilePlus.py* will calculate the gene expression levels based on uniquely mapped reads and randomly select a placement of multi-mapped reads that considers gene expression level.

### 2.5 Running UMI-Reducer across multiple samples

We provide multiple bash scripts to automate running **UMI-Reducer** across multiple samples using a single command. To collapse PCR duplicates across multiple samples, we developed *runCollapsePCRduplicates.sh*. To map multiple samples via tophat2, one can use *runTophat.sh*. To run Genomic Profiler across multiple samples, one can use *runGprofile.sh*.

## 3. Application

We applied **UMI-Reducer** to 22.8 million RNA-Seq reads from a ribosome profiling experiment. RNA-seq libraries from six samples were prepared as described in Section 2.1 and Supplementary Figure S1. Six samples were barcoded during cDNA synthesis and pooled prior to PCR amplification. A single pooled sample was sequenced using the MiSeq Illumina V3 150 cycle kit, which produces reads up to 150nt in length. Applying *extractTag.py* generated an average of 2.7 million reads per sample that contained the full sequence of L32 RNA linker (Supplementary Table S1). Using tophat2 we were able to map 98% of L32-containing reads. Filtering out multi-mapped reads and collapsing technical duplicates resulted in 0.95 million reads per sample available for downstream analyses (36% of original reads per sample). We present the results of **UMI-Reducer** for one of the six samples in Supplementary Figure S2. These results include the number of reads belonging to each genomic category (e.g., coding sequence, UTR, etc.). To estimate ribosome footprint abundance in the original lysate, we report the number of reads that overlap coding regions in each gene.

## 4. Conclusion

We have presented **UMI-Reducer**, a novel computational tool for processing UMI reads and enabling differentiation between PCR and biological duplicates. The proposed approach was coupled with a custom^2^ library preparation protocol to allow accurate estimation of RNA abundance in the original lysate. **UMI-Reducer** can also work as a stand-alone application if UMI reads are in FASTQ format with UMI tags appended to read IDs. The proposed strategy is particularly useful for libraries made from low amounts of starting material, which typically require additional cycles of PCR.

## Supplementary Materials

**Supplementary Figure S1.**
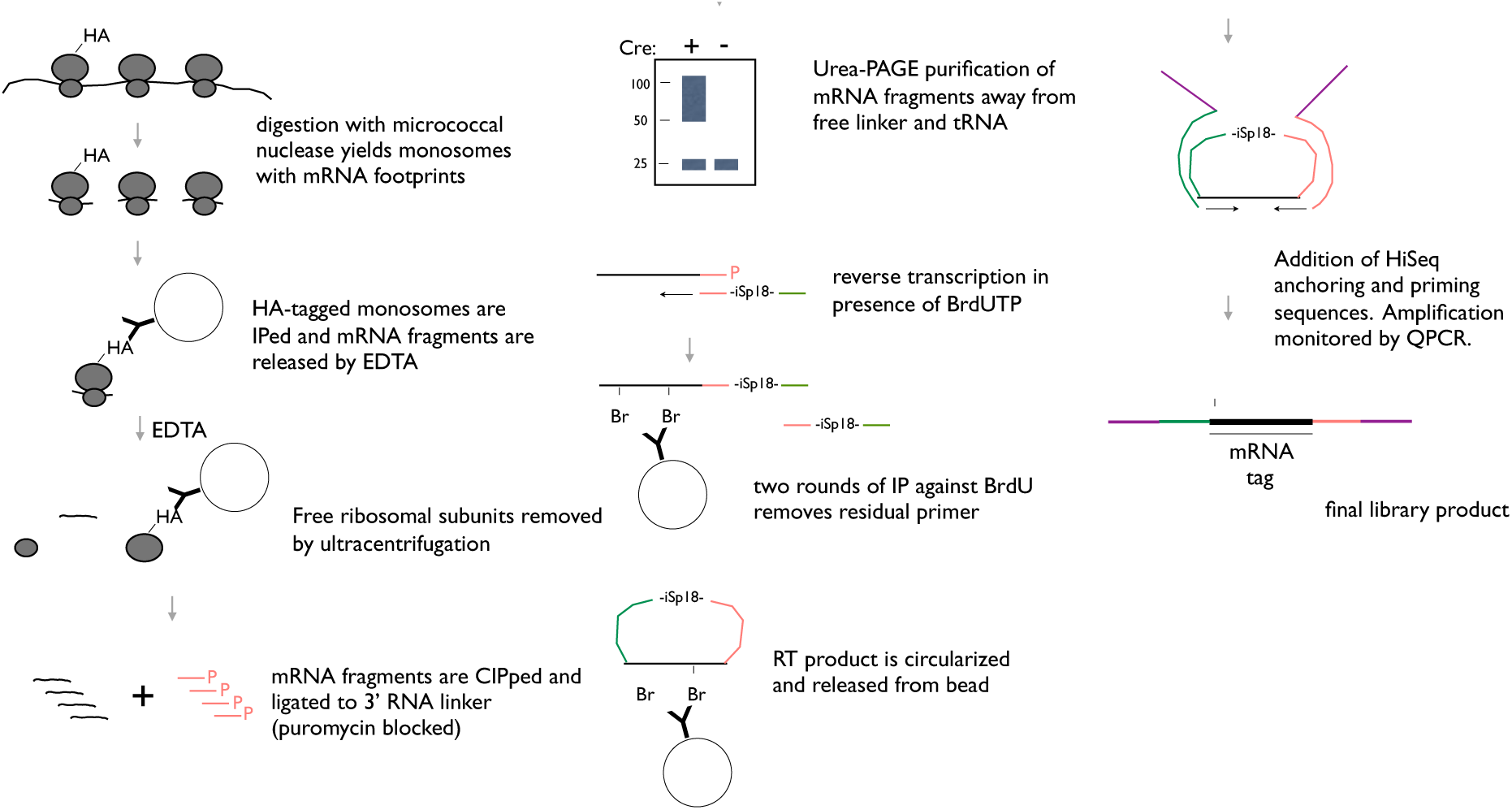
Preparation of ribosome profiling libraries from limited amounts of input material.

**Supplementary Figure S2.**
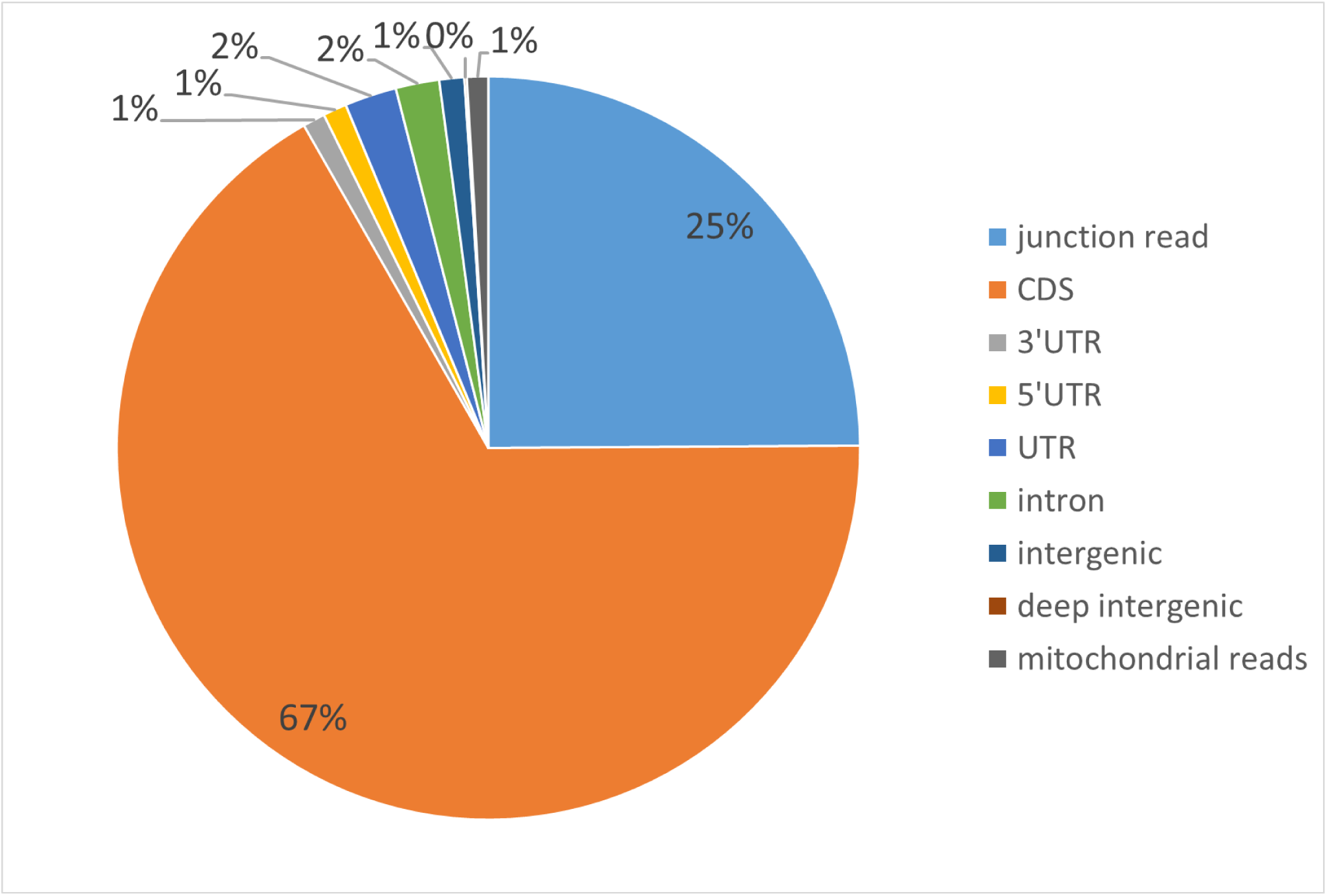
Using the reads available after collapsing PCR duplicates, the genome profiler annotates reads into the following categories: junction reads, CDS, 5’UTR, 3’UTR, intron, intergenic, deep intergenic, and mitochondrial reads (MT). Here, results are shown for one of the six samples listed in Supplementary Table S1.

**Supplementary Table S1.** RNA-seq libraries from the six samples used in this study.

| Sample name | Number of reads | Number of mapped reads | Number of reads after removing multi-mapped reads | Number of reads after collapsing PCR duplicates | Number of mapped reads, % | Number of reads after removing multi-mapped reads, % | Number of reads after collapsing PCR duplicates, % |
| --- | --- | --- | --- | --- | --- | --- | --- |
| SJV076_S1_L001_R1_001_ATCG | 3350230 |  |  |  |  |  |  |
| SJV076_S1_L001_R1_001_CGAT | 3212377 | 3181955 | 2046572 | 1251887 | 99% | 64% | 39% |
| SJV076_S1_L001_R1_001_CTAG.fastq | 1919735 | 1896501 | 1015348 | 618040 | 99% | 53% | 32% |
| SJV076_S1_L001_R1_001_GATC.fastq | 2793065 | 2745509 | 1591371 | 990160 | 98% | 57% | 35% |
| SJV076_S1_L001_R1_001_GCTA.fastq | 3272159 | 3228342 | 2088701 | 1324575 | 99% | 64% | 40% |
| SJV076_S1_L001_R1_001_TAGC.fastq | 1731911 | 1685162 | 963549 | 598462 | 97% | 56% | 35% |

